# A chromosome-length assembly of the Hawaiian Monk seal (*Neomonarchus schauinslandi*) confirms genomic stability in the Pinnipeds and a prolonged history of “genetic purging”

**DOI:** 10.1101/2022.03.31.486393

**Authors:** David W. Mohr, Stephen J. Gaughran, Justin Paschall, Ahmed Naguib, Andy Wing Chun Pang, Olga Dudchenko, Erez Lieberman Aiden, Deanna M. Church, Alan F. Scott

## Abstract

The Hawaiian monk seal (HMS) is the single extant species of tropical earless seals of the genus *Neomonachus*. The species survived a severe bottleneck in the late 19^th^ century and experienced subsequent population declines until becoming the subject of a NOAA-led species recovery effort beginning in 1976 when the population was fewer than 1000 animals. Like other recovering species, the Hawaiian monk seal has been reported to have reduced genetic heterogeneity due to the bottleneck and subsequent inbreeding. Here we report a chromosomal reference assembly for a male animal produced using a variety of methods including linked-read sequencing, Hi-C contiguity mapping, optical genome mapping, and nanopore long read sequencing. The final assembly consisted of 16 autosomes, an X and portions of the Y chromosomes. We compared variants in the reference animals to nine other HMS and to the human reference NA12878 confirming a low level of variation within the species and one-eighth that of the human reference. A lack of variation in several MHC genes was documented suggesting that this species may be at risk for infectious disease. Lastly, the HMS chromosomal assembly confirmed significant synteny with other pinnipeds. This reference should be a useful tool for long-term management of HMS and evolutionary studies of other carnivorans.

## INTRODUCTION

High quality non-human genomes, especially mammalian genomes, are needed to 1) better constrain the limits of naturally occurring nucleotide variation, 2) identify conserved protein and regulatory regions that may explain the distinct morphological or physiological characteristics of species, 3) establish the correct relationship of DNA variants to genes for association studies, 4) improve our understanding of evolutionary relatedness and mechanisms, and 5) aid efforts for species conservation and management. The genome of the Hawaiian monk seal (*Neomonarchus schauinslandi*) is especially relevant as it underwent a severe bottle-neck due to overhunting in the 19^th^ century and the population continued to fall throughout the 20^th^ century. Its sister species, the Caribbean monk seal (*N. tropicalis*) was last observed in 1952 and declared extinct in 2008. With the passing of the Endangered Species Act in 1973 the National Oceanic and Atmospheric Administration (NOAA) listed the Hawaiian monk seal as endangered in 1976 and published species recovery plans in 1993 and 2007 (ref: Recovery plan for the Hawaiian monk seal *Monachus schauinslandi*^1^, https://repository.library.noaa.gov/view/noaa/3521). Studies of genetic heterogeneity for this species have demonstrated a significant loss of heterozygosity as a result of the bottleneck and the presumed inbreeding (e.g., Schultz et al, 2009). Previously, we reported a scaffold-level assembly (Mohr et al., 2017), which we have improved upon to produce a chromosome-length genome that will serve as an accurate reference for *N. schauinslandi*. We have compared this reference to that of other HMS and to selected human samples to assess overall heterogeneity, especially in genes that may be relevant to risk of disease.

The genome was assembled from a single male animal, RE74 aka “Benny,” who is well-known on Oahu, and has been followed since his birth on Kauai in 2002. RE74 has been captured on various occasions for veterinary care due to injuries including hook ingestion (Fig 1). On one such occasion blood was taken, DNA isolated and a variety of methods was used to assemble a chromosomal-length genome assembly which confirmed a karyotype of 16 autosomes, along with the X chromosome and portions of the Y. The genome was annotated by NCBI and measures of genome quality and completeness were assessed. Using the linked-read data we were also able to create phased haplotype blocks for the autosomes, compare heterozygosity at specific loci related to disease susceptibility relative to the human gold-standard, NA12878, and to nine additional HMS. We also compared RE74 assembly to those of other seal genomes available through the DNAzoo.org and at NCBI to address the question of syntenic conservation.

**Fig. 1.**
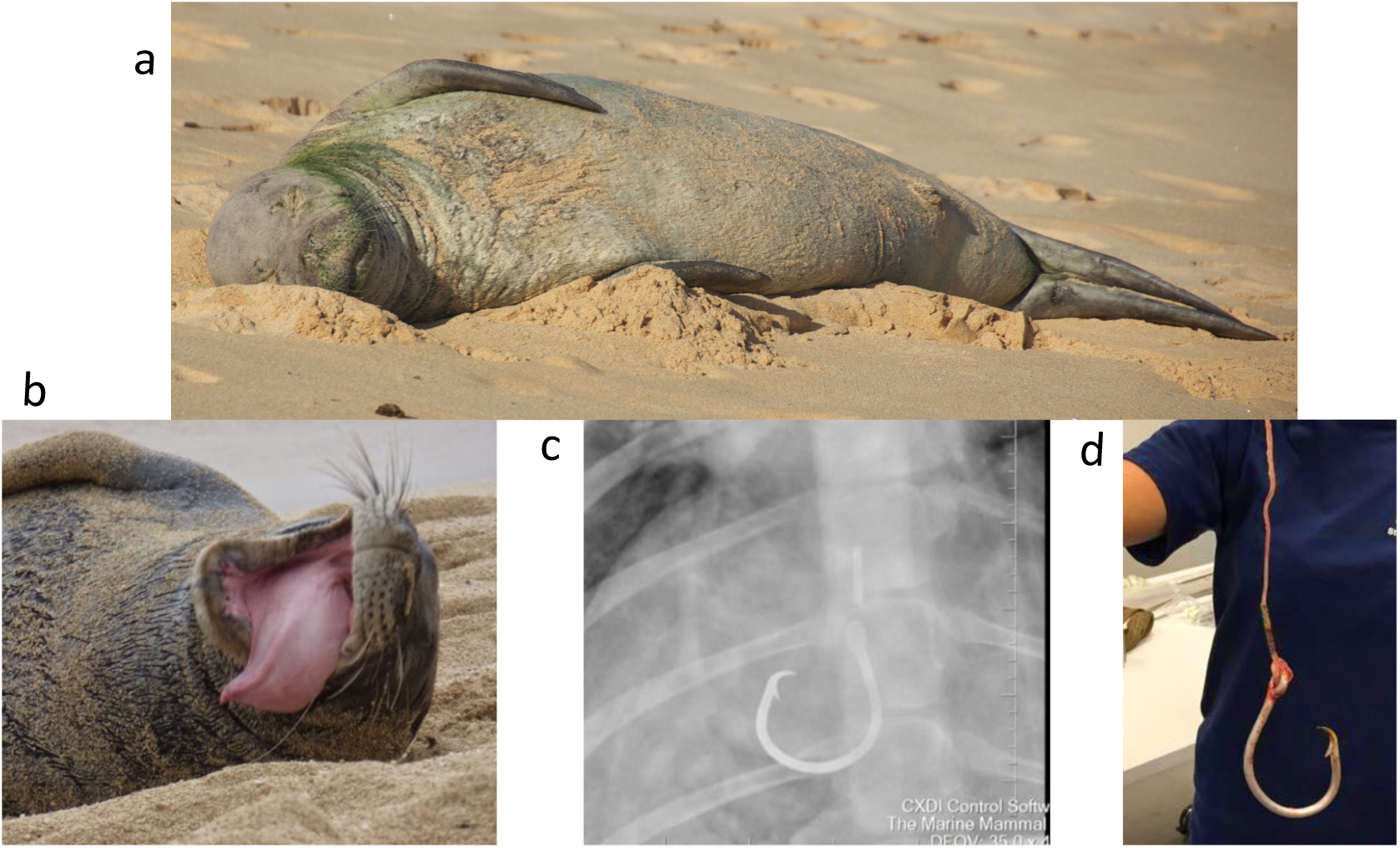
Montage of “Benny” (RE74) a. Asleep on east Oahu in 2009, b. In distress after swallowing a fishing hook and line (2015), c. X-ray prior to surgery and d. post-surgery. Images courtesy of NOAA

## METHODS

### Sample

A male animal was chosen so that both sex chromosomes would be available for study and that assembly of the X-chromosome would not be confounded by heterozygosity. The sample was collected under the Marine Mammal Health and Stranding Response Program Permit No. 932-1905-00/MA-009526 in August 2015. Blood (∼10 ml) was collected in EDTA vacutainers, shipped to Baltimore and processed within two days of collection. The viability of the cells was assessed by trypan blue exclusion. 1 ml of whole blood was stored in LN_2_ while 9 ml (∼1.85×10^6^ cells/ml) were used for lymphocyte separation and subsequent DNA isolation for optical genome mapping or Hi-C library preparation. As for other non-captive animals, sample collection was opportunistic and limited which restricted our use of certain technologies. The overall sequencing strategy is shown in Supplement Table 1 and evolved as technology and analysis tools improved.

### Linked-read sequencing

DNA was isolated using MagAttract (Qiagen) and the molecular weight assayed by pulsed field gel electrophoresis. High molecular weight (HMW) gDNA concentration was quantitated using a Qubit Fluorometer, diluted to 1.25 ng/ul in TE buffer, and denatured following manufacturers recommendations. Denatured gDNA was added to the reaction master mix and combined with the Chromium bead-attached primer library and emulsification oil on a Chromium Genome Chip (10X Genomics). Library preparation was completed following the manufacturer’s protocol (Chromium Genome v1, PN-120229). Sequencing-ready libraries were quantified by qPCR (KAPA) and their sizes assayed by Bioanalyzer (Agilent) electrophoresis. The linked-read library was sequenced (151 × 9 × 151) using two HiSeq 2500 Rapid flow cells to generate 975 M reads with a mean read length of 139 bp after trimming. The read 2 Q30 was 87.93% and the weighted mean molecule size was calculated as 92.33 kb. Mean read depth was ∼61X. The sequence was analyzed using Supernova software (10X Genomics; Weisenfeld et al., 2016) which demultiplexed the Chromium molecular indexes, converted the sequences to fastq files and built a graph-based assembly. The assemblies, which diverge at “megabubbles,” consisted of two “pseudohaplotypes.” The sequence data were originally analyzed using Supernova 1.0 and then repeated using v1.1 which estimates gap sizes rather than introducing an arbitrary value of 100 Ns. As noted above, the Supernova scaffolds were used by the Bionano Hybrid Scaffold tool to create sequence assemblies. 10X linked reads. Subsequently, we reanalyzed the linked-read data with Supernova version 2.1.1 on 1260.32M reads and ∼78X read depth. Supernova 2.1.1 increased scaffold length N50 from ∼22 Mb to ∼62 Mb. [Supplemental Fig. 1].

### Nanopore sequencing

We used Oxford Nanopore sequencing to obtain long reads for gap filling. DNA was isolated using the agarose plug method (www.bionanogenomics.com) or with the Circulomics nanobind method (www.circulomics.com). Libraries were made using either the rapid transposase (RAD-004) or ligation methods (LSK-110) and run on a GridION. A total of 4.26M reads were obtained with molecule N50s of approximately 32 kb producing about 7.3X total read depth. PILON v1.22 was run in gap mode for filling.

### Bionano Genomics

Two separate optical genome maps were created for RE74. The second map was done to take advantage of improved chemistry and instrumentation.

Sample 1: Optical genome mapping of large DNA (Xiao et al., 2007) incorporates fluorescent nucleotides at sequence specific sites, visualizes the labeled molecules and aligns these to each other and to a DNA scaffold (Shelton et al., 2015). Lymphocytes were processed following the IrysPrep Kit for human blood with minor modifications. Briefly, PBMCs were spun and resuspended in Cell Suspension Buffer and embedded in 0.6% agarose (plug lysis kit, BioRad). The agarose plugs were treated with Puregene Proteinase K (Qiagen) in lysis buffer (Bionano Genomics) overnight at 50°C and shipped for subsequent processing (S. Brown, KSU). High Molecular Weight (HMW) DNA was recovered by treating the plugs with Gelase (Epicenter), followed by drop dialysis to remove simple carbohydrates. HMW DNA was treated with Nt. BspQI nicking endonuclease (New England Biolabs) and fluorescent nucleotides incorporated by nick translation (IrysPrep Labeling-NLRS protocol, Bionano). Labeled DNA was imaged on the Irys platform (Bionano) and more than 234,000 Mb of image data were collected with a minimum molecule length of 150 kb. Alignment was done with Bionano Solve3.2.2_08222018 software.

Sample 2: Optical genome mapping was repeated using the DLE-1 label and the Saphyr instrument. DNA from approximately 5M lymphocytes were isolated, labeled and imaged at Bionano Genomics. Alignment to the post-Hi-C assembly was performed using Bionano Solve v3.6. As with sample 1, molecules smaller than 150 kb were excluded. The second optical genome maps were aligned to the predicted assembly resulting from the Hi-C assembly. The maps were reviewed using the online Bionano Access viewer Raw data, along with hybrid assembly files, are available at NCBI (SUB11144351)

### Arima Hi-C and 3d-dna

Approximately 1M lymphocytes were sent to Arima Genomics for Hi-C library prep. The library was sequenced locally on a NovaSeq 6000 to a read depth of 60.8X. The Hi-C data were aligned to the linked-read Supernova scaffolds. Hi-C genome assembly was performed using the 3D-DNA pipeline (3d-dna v180922) (Dudchenko et al. 2017) and the output was reviewed using Juicebox Assembly Tools (Dudchenko et al. 2018). The Hi-C data are available on www.dnazoo.org/assemblies/Neomonachus_schauinslandi, where they were visualized using Juicebox.js, a cloud-based visualization system for Hi-C data (Robinson et al. 2018). Genome assembly before and after the Hi-C scaffolding is shown in Supp. Fig. 2.

**Fig. 2.**
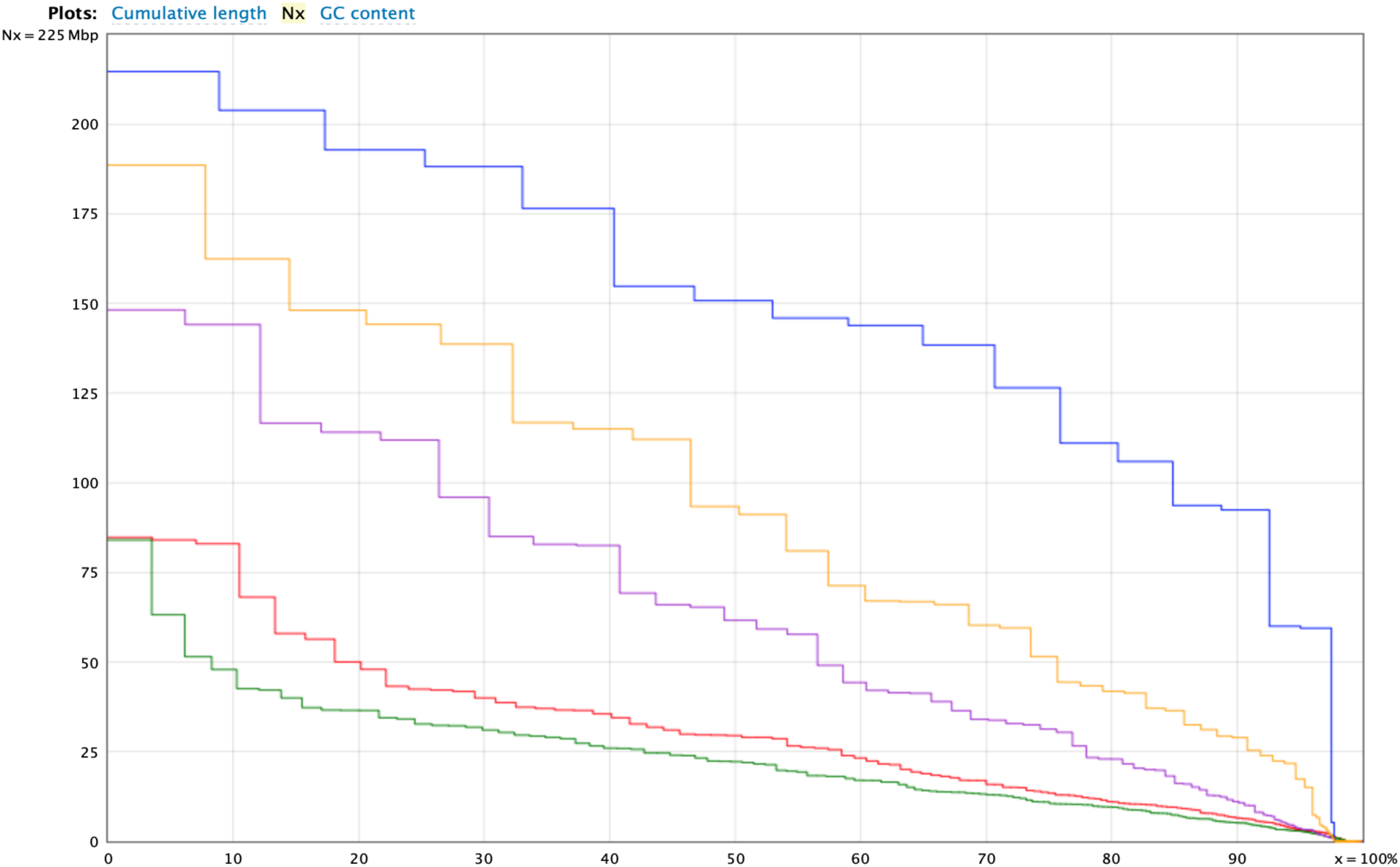
QUAST plot showing the improvement of assembly metrics from version 1 (NCBI ASM220157v1) to version 2 (NCBI ASM220157v2) and the small contribution of unplaced scaffolds in v2. The green line represents scaffolds remaining after the second optical genome mapping.

### Final Editing

The near final assembly from the methods described above was manually edited with respect to the second optical genome map (Bionano Access v1.6.1). This editing mainly adjusted inverted sequence within scaffolds and altered the length of N blocks introduced by linked-read sequencing assembly. We did not attempt to remove large N blocks which were generated largely by the Bionano software resolving our Hi-C scaffolds with the optical genome maps. Manual inspection of the optical genome maps aligned with the sequence frequently showed that the spacing of DLE-1 sites in blocks of long Ns corresponded to repetitive sequence. The final edited version was submitted to NCBI for annotation.

### Haplotype analysis

The linked read sequence allowed phasing of sequences sharing indexes into haplotype blocks. We used short read data aligned to the final assembly to run Long Ranger 2.2.2. Loupe ver 2.1.2 (2.4) was used to visualize the pseudohaplotypes. A loupe file for NA12878 was available at 10X Genomics.

### Quality Assessments

QUAST (Gurevich et al., 2013) was used to generate N50 plots and comparative metrics for the assembly methods. BUSCOv5 (Benchmarking Universal Single Copy Orthologs), a tool for assessing genome completeness (Simão et al., 2015), was run on the final assembly.

### Conserved Synteny Analysis

We compared the HMS reference to the domestic dog, cat, and other seal assemblies using minimap2 (Alonge et al, 2019) and plotted these with D-GENIES (Cabanettes & Klopp, 2018).

### Annotation and QC

Annotation of the version 1 and 2 scaffolds (ASM220157v1/v2) was performed at NCBI using the Eukaryotic Genome Annotation Pipeline (https://www.ncbi.nlm.nih.gov/genome/annotation_euk/process/). We used BUSCO ver5 to identify shared genes predicted to be present or missing in our chromosomal assembly vs. those of other species characterized by the minimap2 alignments.

### Short read sequencing and variant calling

For the additional 9 HMS, standard Illumina short-read libraries were prepared from flipper-punch DNA, indexed and sequenced. Alignment, variant calling, and quality control were accomplished using the DRAGEN Germline v3.7.5 pipeline on the Illumina BaseSpace Sequence Hub platform (Zhao et al 2020). FASTQ files were aligned to the RE74 reference (ASM220157v2), producing CRAM files with duplicate reads flagged. Quality control statistics for alignment and variant calling were downloaded using the Basespace CLI, and analyzed using in-house aggregation and visualization pipelines. No significant human cross sample contamination was observed using the DRAGEN contamination detection tool. Once adequate coverage and data quality were confirmed, joint variant calling was performed, producing a multi-sample VCF file. Filter flags were applied using the DRAGEN Germline default hard-filter quality control parameters, flagging variants which do not meet the criteria of (QUAL score < 10.41 SNV or QUAL score < 7.83 for indels) or (lod_fstar > 6.3). Systematic base calling errors are corrected by default using the Base Quality Drop off algorithm.

### Demographic reconstruction

We modeled the demographic history of the species using the sequentially Markhovian coalescent as implemented in MSMC2 (Schiffels and Durbin 2014, Schiffels and Wang 2020). We followed the workflow provided by Schiffels and Wang 2020. This included generating a mask file for the reference genome to exclude unmappable regions, and individual mask files for each genome to exclude regions of unexpectedly high or low coverage. We then ran the unphased data of four medium-to-high coverage samples through MSMC2, with a per-generation mutation rate of 7.0 × 10^−9^ (Peart et al. 2020) and a generation time of 13 years (Schulz et al. 2010).

## RESULTS

The completed assembly consisted of 16 autosomes, the X chromosome, portions of the Y chromosome and the mitochondrial genome. A QUAST plot (Fig. 2) shows the improvement in scaffold contiguity between NCBI ASM220157v1 and v2. Changes in the NCBI annotation metrics are shown in Supp. Table 3. Improvements may reflect both changes in the contiguity of the assembly as well as evolution of the assembly pipeline. The genome report for ASM220157v2 is available at https://www.ncbi.nlm.nih.gov/assembly/GCF_002201575.2/. Key metrics comparing the two assemblies are shown in Supplement Table 3.

Although imperfect, comparison of genes expected to be conserved is a useful tool for comparing the completeness of genomes. BUSCO (Simão, et al 2015), a collection of predicted conserved genes in different taxa, was used on our HMS chromosome-length assembly together with the small number of unplaced scaffolds. BUSCO genes can differ between taxa if a genome is incomplete or has errors that make gene prediction inaccurate. Differences between taxa can also be the result of selection allowing genes to be gained or lost over evolutionary time. BUSCO metrics for RE74 are comparable to other genomes (Table 1) suggesting both completeness of this assembly as well as that of the other species examined.

**Table 1.** BUSCO comparisons (9226 BUSCO gene groups) of HMS to other pinniped genomes. Ma, *Mirounga angustirostris* (N. Elephant seal), Pv, *Phoca vitulina* (Harbor seal), Hg, *Halichoerus grypus* (Gray seal), Zc, *Zalophus californianus* (California sea lion), Or, *Odobenus rosmarus* (Walrus), Clf, *Canis lupus familiaris* (Domestic dog), Fc, *Felis cattus* (Domestic cat), Hs, *Homo sapiens* (GRCh38).

|  | RE74 | Ma | Pv | Hg | Zc | Or | Clf | Fc | Hs |
| --- | --- | --- | --- | --- | --- | --- | --- | --- | --- |
| Complete | 8845 | 8712 | 8849 | 8542 | 8878 | 8875 | 8742 | 8836 | 8856 |
| Comp & Single-Copy | 8617 | 8467 | 8651 | 8352 | 8543 | 8599 | 8560 | 8761 | 8450 |
| Comp & Duplicated | 228 | 245 | 198 | 190 | 335 | 276 | 182 | 75 | 406 |
| Fragmented | 100 | 175 | 90 | 242 | 83 | 82 | 152 | 111 | 127 |
| Missing | 281 | 339 | 287 | 442 | 265 | 269 | 332 | 279 | 243 |

We noted that there was a pattern of shared BUSCO “missing” genes between selected species (Table 2) that followed their expected phylogenetic distance as calculated from timetree.org estimates of divergence time. We interpret this result to mean that many of the missing genes are truly absent in these related taxa and reflect a shared history or shared biology.

**Table 2.** Number of shared “missing” BUSCO genes between taxa. TT est. is the estimate divergence time from HMS from Timetree.org.

| Total missing |  | HMSRE74 | Ma | Pv | Hg | Zc | Or | Clf | Fc | Hs |
| --- | --- | --- | --- | --- | --- | --- | --- | --- | --- | --- |
|  | **TT est.** | - | **13.5** | **18.4** | **18.4** | **26** | **26** | **46** | **54** | **96** |
| 281 | HMS | - | 188 | 185 | 179 | 164 | 167 | 129 | 128 | 99 |
| 339 | Ma |  | - | 186 | 185 | 153 | 155 | 127 | 133 | 100 |
| 332 | Pv |  |  | - | 207 | 172 | 170 | 131 | 135 | 111 |
| 332 | Hg |  |  |  | - | 160 | 166 | 125 | 133 | 108 |
| 265 | Zc |  |  |  |  | - | 184 | 133 | 126 | 106 |
| 269 | Or |  |  |  |  |  | - | 127 | 134 | 103 |
| 332 | Clf |  |  |  |  |  |  | - | 122 | 104 |
| 279 | Fc |  |  |  |  |  |  |  | - | 106 |
| 243 | Hs |  |  |  |  |  |  |  |  | - |

Arnason (1974) observed that pinniped chromosomes show “pronounced karyotypic uniformity” and have either 15 or 16 autosomes (Beklemisheva, et al., 2020). Our earlier methods could not definitively resolve the correct number of HMS chromosomes but the addition of Hi-C confirmed 16 autosomes. We renumbered all Hi-C autosomes according to their length and identified the X and Y by synteny. We compared the HMS chromosomal assembly to those of other seals and carnivores using minimap2 and plotted these with D-GENIES (Cabanettes & Klopp, 2018) as shown in Fig. 3. Fragmentation or rearrangement of syntenic blocks in such an analysis can occur either by errors in the assemblies or true chromosomal events. Assuming correct assemblies, the seal DG plots confirm conservation between species reflecting phylogenetic relatedness.

**Fig. 3.**
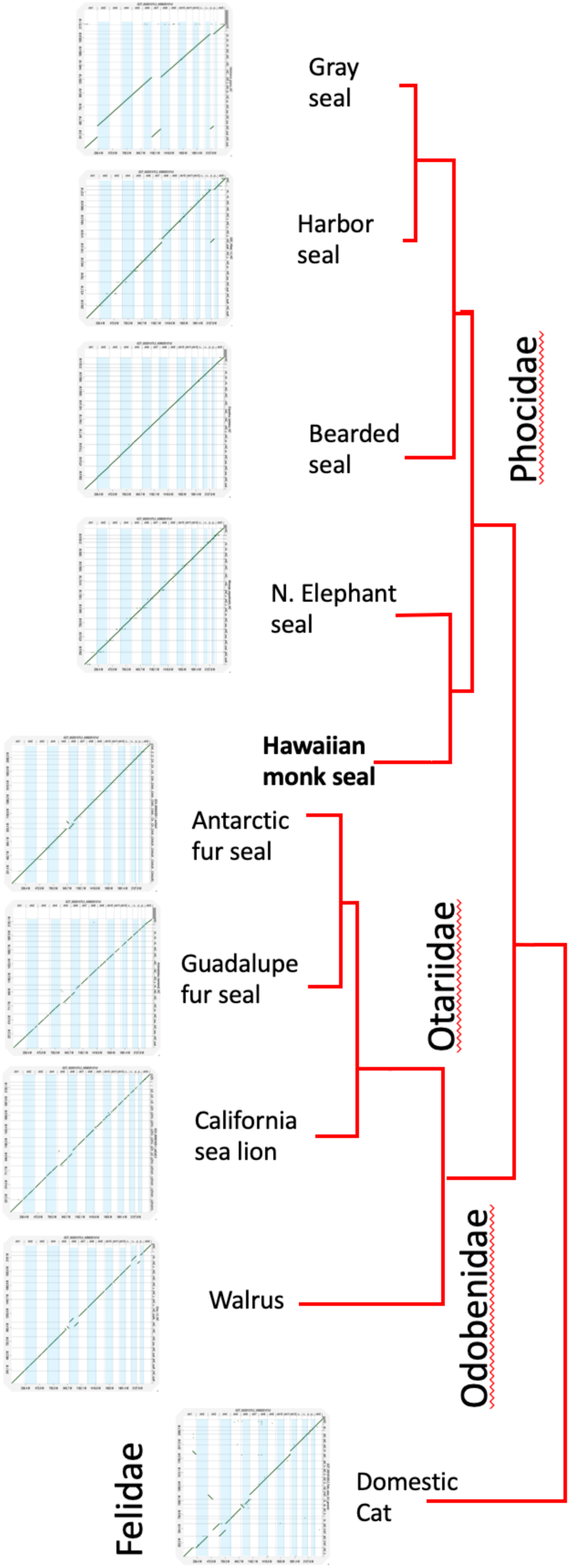
D-GENIES (Cabanettes & Klopp, 2018) plots of HMS aligned to other pinnipeds for which chromosome-length assemblies are available as well as the domestic cat. Tree lengths are redrawn from REF. The specific genomes compared are listed in Supp. Table2. Because chromosome orientations are arbitrary based on their order in NCBI we reverse-complemented chromosomes in some species prior to alignment so that the chromosomes are in the same direction relative to HMS. Enlarged individual alignments are shown in Supplement Fig. 3. D-GENIES filtering was adjusted individually for each plot to maximize sequence identity but minimize small matches.

D-GENIES plots showed near perfect chromosome to chromosome synteny between HMS and the Northern Elephant seal (*Mirounga angustirostris*) the taxonomically closest species for which an assembly is available as well as the Bearded seal (*Erignathus barbatus*) both of which have 16 autosome pairs as does the Hawaiian monk seal. The Gray seal (*Halichoerus grypus*), another phocid, has a fusion of HMS chromosomes 7 and 15. The harbor seal (*Phoca vitulina)*, also has one fewer autosome pair due to the same fusion. In all of the phocids the remaining autosomes are syntenic to HMS. In the more distant Guadalupe fur seal (*Arctocephalus townsendi)* with 17 autosome pairs, chromosomes 8 and 10 share synteny with HMS 5 and chromosomes 10 and 15 share with HMS 6. The same is true for the Antarctic fur seal (*Arctocephalus gazella)*. The most distant of the pinnipeds, the walrus (*Odobenus rosmarus)* with 15 autosome pairs, is the most distinct with HMS chromosome 5 syntenic to portions of Oros chromosome 16 and 13, HMS 6 is syntenic to Oros 13 and portions of 3, HMS 7 also matches portions of Oros 3, and HMS 13 and 15 each match portions of Oros 15.

Among the two other carnivores we compared, domestic dog and cat, the cat is most similar karyotypically despite being taxonomically more distant. Cats have 18 autosome pairs and, as shown in supplement Fig. 3, HMS 1 is the equivalent of cat 018731.3 and parts of 018724.3, HMS 3 is syntenic with parts of chr C1 (NC_018730.3) and chr A1 (NC_018723.3), HMS 4 with cat chr F2 (NC_018740.3) and part of chr C1 (NC_018730.3), HMS 5 with chr B4 (NC_018729.3) and chr E3 (NC_018738.3), HMS 7 with part of chr A1 (NC_018723.3), and HMS 12 with part of chr A2 (NC_018724.3). The dog is the most different with 38 autosome pairs with at least 68 syntenic blocks that are fragmented, rearranged or inverted relative to the monk seal (Supplement Fig. 3). As expected, only the X chromosome is universally conserved.

As with other mammals the X chromosome is, as expected, highly syntenic. The Y chromosome was the most difficult to assemble in part because of highly repetitive sequence that accumulates variation rapidly over evolutionary time and the similarity of genes in the pseudoautosomal regions that can be mistaken as from the X. Hi-C analysis identified three scaffolds thought to be from the Y which we assembled manually using the California Sea Lion assembly (CM019820.2; Peart et al 2021) and other mammals as guides. Approximately 2.59 Mb of our assembly includes genes identified on the California Sea Lion although the order of some regions is different. Whether this represents errors in our assembly or true differences remains unresolved. Table 3 lists 20 Y-specific genes that are shared with *Zalophus*, the only other seal for which Y chromosome sequence was available.

**Table 3.** List of Y chromosome genes identified in RE74 from NCBI annotation with homologous sequences in the *Zalophus californianus* Y (CM019820.2).

| Position/Mb | Locus ID | RNA Sequence | NCBI annotation |
| --- | --- | --- | --- |
| 2.66 | LOC110582119 | XM_021692074.1 | ubiquitin-like modifier-activating enzyme 1 |
| 2.64 | LOC110582047 | XM_021691967.1 | BCL-6 corepressor-like |
| 2.89 | LOC110581036 | XM_021690741.1 | gamma-taxilin-like |
| 2.95 | SRY | XM_021693078.1 | sex-determining region Y, SRY |
| 3.05 | LOC110582200 | XM_021692141.1 | cullin-4B-like |
| 3.33 | LOC110591777 | XM_021702682.1 | eukaryotic translation initiation factor 1A, X-chromosomal |
| 3.42 | LOC110591779 | XM_021702683.1 | heat shock transcription factor, Y-linked-like |
| 3.50 | LOC110591787 | XM_021702687.1 | lysine-specific demethylase 5D |
| 3.56 | LOC110591776 | XM_021702680.1 | zinc finger X-chromosomal protein-like |
| 3.66 | LOC110591775 | XM_021702679.1 | eukaryotic translation initiation factor 2 subunit 3, Y-linked-like |
| 4.41 | OFD1 | XM_021692108.1 | OFD1, centriole and centriolar satellite protein (OFD1), partial mRNA |
| 4.46 | LOC110580574 | XM_021690271.1 | histone demethylase UTY-like |
| 4.66 | LOC110581163 | XM_021690923.1 | ATP-dependent RNA helicase DDX3X-like |
| 4.70 | LOC110581157 | XM_021690911.1 | probable ubiquitin carboxyl-terminal hydrolase FAF-X |
| 4.81 | LOC110580811 | XM_021690464.1 | RNA-binding motif protein, X chromosome-like |
| 4.83 | LOC110580801 | XM_021690458.1 | probable ubiquitin carboxyl-terminal hydrolase FAF-Y |
| 4.89 | LOC110580822 | XM_021690474.1 | serine protease 55-like |
| 4.91 | LOC110580830 | XM_021690484.1 | serine protease 52-like |
| 5.17 | LOC110581101 | XM_021690826.1 | thymosin beta-4-like |
| 5.25 | LOC110584133 | XM_021694172.1 | ras-related protein Rab-9A-like |

### Heterozygosity estimates and phasing

We created a multi-sample hard-filtered VCF file using reads for RE74 and for the additional nine HMS using GATK as implemented on the Illumina DRAGEN platform and aligned to the chromosomal assembly. Using VCFtools we compared the RE74 SNPs to autosomal SNPs from NA12878 and noted an overall reduction of heterozygosity in RE74 that was approximately one-eighth that of the human reference (350,812/3,998486=12.4%). Given the smaller size of the combined seal autosomes (∼2.23 Gb) vs. NA12878 (∼3.09 Gb) the SNPs per Mb average ∼157 for RE74 vs 1,262 for NA12878. Considering the caveats in making such comparisons these numbers agree with the empirical data we observed from gene-to-gene comparisons visualized in Loupe and IGV. Long Ranger (10X Genomics, v2.2.2) statistics for RE74 indicated that 96% of SNPs were phased, the phase block N50 was 557.7 kb and the longest phase block was 7.27 Mb. We compared phase blocks between RE74 and NA12878 in the Loupe viewer. A representative example of a 1.28 Mb phase block including the KCNAB1 and PLCH1 genes is shown in Supp. Fig. 4.

### MHC genes

Immunity-related genes have been previously studied in the Hawaiian monk seal and their lack of heterogeneity documented (de Sá et al., 2019). Fig. 4 compares heterozygosity in NA12878 for HLA-DQA1 and HLA-DQB1 to the orthologous HMS MHC genes. The same observation was made for HLA DMB and DMA (Fig. 5a) and HLA-DOA and HLA-DOB (Fig. 5b) where we compared RE74 and 10 other seals and to four human CEPH samples.

**Fig 4.**
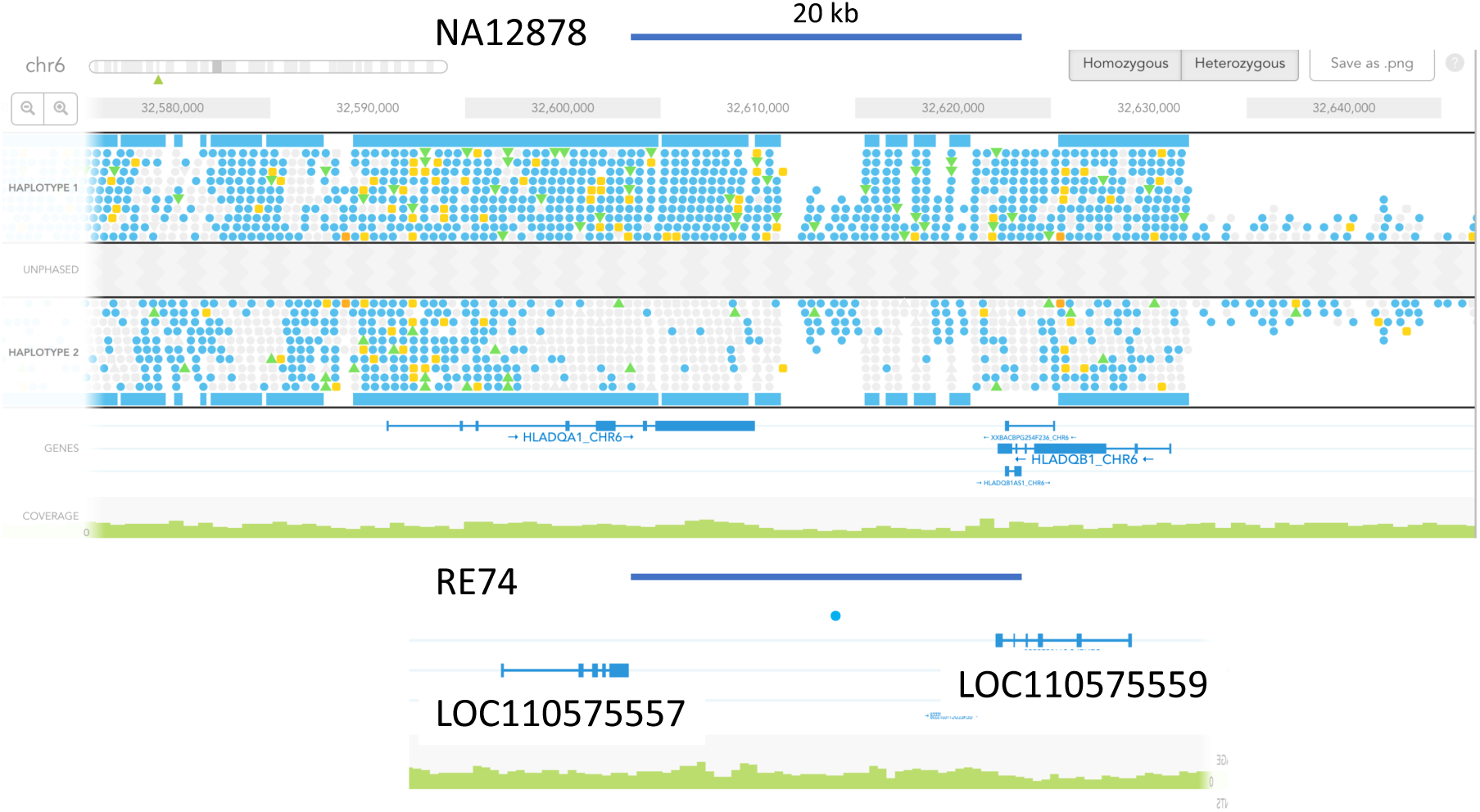
Absence of heterozygosity in RE74 at loci orthologous to human DQA1 and B1. Comparison of phased variants in NA12878 HLA DQA1 and B1 genes visualized in the Loupe viewer (10X genomics) compared to orthologous RE74 genes (de Sá, et al 2019), LOC110575557 an LOC110575559. The figures were adjusted to the same scale (the blue bar represents 20 kb). Blue dots represent SNPs and yellow dots are indels. Blue squares represent multiple SNPs that are not resolved at this scale. One low quality intergenic SNV was observed in RE74.

**Fig 5a.**
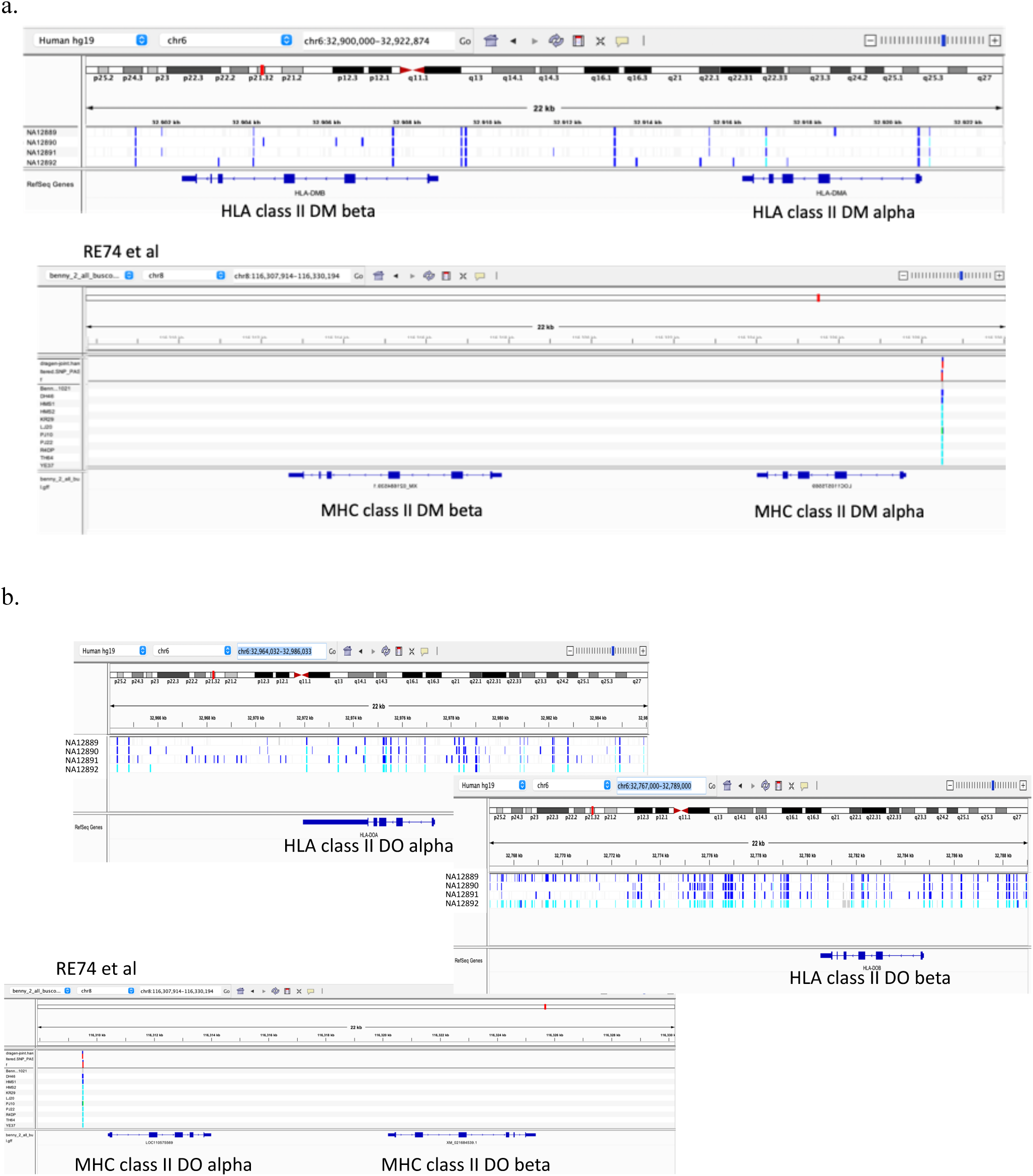
Low frequency of heterozygous positions flanking the MHC class II loci DM alpha, DM beta, DO alpha and DO beta in RE74 and nine other HMS genomes as visualized in IGV. Human genomes are CEPH NA12889-12892. Seal SNVs occur at only one position for all animals for both regions. 5b. Comparisons of NA12889-12892 for HLA DO alpha and DO beta (the human genes are split due to their larger intergenic sequence). Seal SNVs occur at a single position for all 10 animals for both regions. Panel width is 22 kb for all panels.

A Multiple Sequentially Markovian Coalescence (MSMC) analysis (Schiffels and Durbin, 2014) was performed for four individual HMS (Fig. 6) and illustrates a notable decline in effective population size over time, starting around 100,000 years ago. The analysis is consistent with a Hawaiian monk seal population that has been relatively small size (Ne<2000) for tens of thousands of years.

**Fig. 6.**
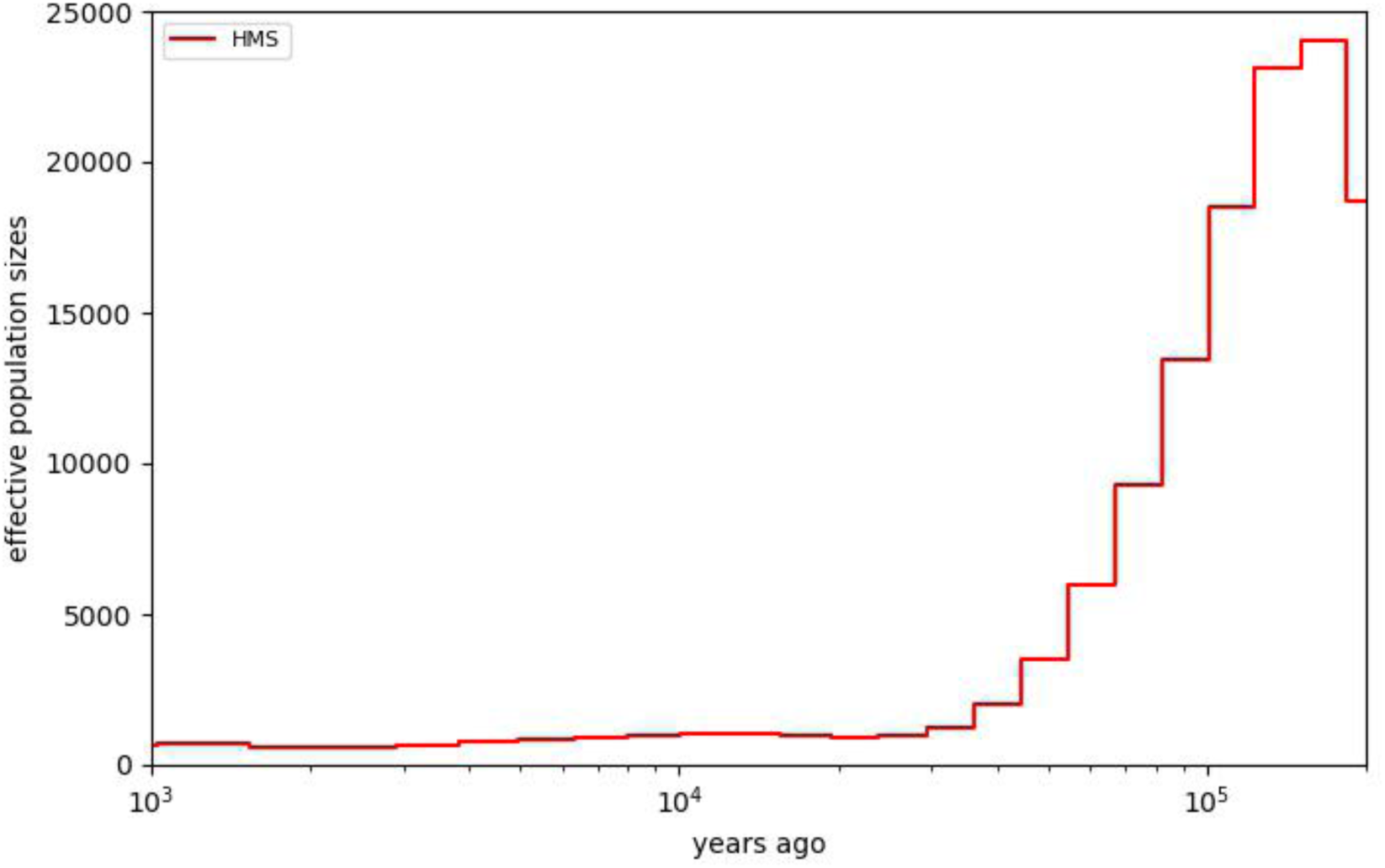
MSMC plot showing small and declining Ne over the last 200,000 years.

## DISCUSSION

Comparative genomics is producing an unprecedented amount of sequence useful to explore fundamental questions about evolutionary biology and population genetics, among others. Comparative sequence is also increasingly useful in human studies such as improving predictions of deleterious mutations (e.g., Frazer et al 2021, Sundaram et al, 2018). The recent development of long DNA methods has allowed high-quality assemblies to be produced more quickly and with greater accuracy. This has led to many independent efforts to characterize a broad cross-section of living plants and animals. In this study we applied a variety of technologies and analysis tools to characterize an endangered species that is recovering from a population that may have ebbed with an effective population size of as few as 25 individuals (Shultz, et al 2009). In this study we estimate that “Benny” has ∼12.4% percent of the heterozygosity seen in one well-characterized human, a number which is supported when compared to several additional HMS. The reduced heterozygosity is comparable to that observed in archaic humans which have been attributed to first cousin or half-sib mating (Prüfer, et al, 2014). The marked loss of heterozygosity was particularly notable in orthologous MHC genes where we observed virtually no variants in HMS relative to CEPH genomes. The consequences of reduced MHC class II variability remain to be determined but an expectation of greater susceptibility to infectious disease may be one. A secondary consequence of the markedly reduced heterozygosity in RE74 is that phase blocks were generally smaller than in NA12878. This was likely the consequence of insufficient variation available to combine linked reads into phased scaffolds. If so, then phasing of other genomes with markedly reduced heterozygosity may be difficult.

Population bottlenecks can result in a high incidence of genetic disorders due to homozygosity of deleterious recessive alleles. However, as with artificial selection, purging of rare recessives can occur relatively quickly in small populations. In contrast, large populations that have experienced a rapid decline would be expected to show residual variation that was present prior to the bottleneck. In the case of Hawaiian monk seals, it seems that allelic purging for a prolonged period, and as indicated by the MSMC analysis, is the best explanation for the extremely low heterozygosity that we and others have observed and the fact that pups with obvious genetic disorders are seldom reported. While the apparent loss of deleterious recessives may be advantageous the loss of an allelic “reserve” could make the species more susceptible to infectious disease or environmental change (e.g., Kardos et al 2021). Because the demographic analysis suggests historically low populations prior to human activity this could be a consequence of the lower productivity of tropical oceans and climatic cycles such as El Niño, that may limit resource availability.

As noted above, BUSCO analysis confirmed that all the seal genomes compared in this study were of comparable quality in terms of completeness and they shared both similar and common “missing” BUSCOs. The loss of genes in particular taxa is a well-documented phenomenon. Whether the missing BUSCOs we observed in pinnipeds merely reflect a shared history or are biologically significant will require future study but multiple instances of gene loss as adaptations to aquatic life have been documented in cetaceans and hippos (Springer et al., 2021).

All the pinnipeds for which chromosomal-length assemblies are available have well conserved synteny. Our D-GENIES comparisons of RE74 to other seals are in agreement with their relatedness both in terms of chromosome counts and overall synteny and supports previous cytogenetics studies (e.g., Beklemisheva et al, 2020). The genome of the domestic cat also shows similarity to the pinnipeds with several autosomes completely syntenic suggesting that seals and felids may share a conserved genome architecture with early carnivorans. In contrast, the canines that are taxonomically more closely related to seals have undergone considerable change at the chromosomal level (e.g., Becker et al, 2011). The consequences of the high rate of karyotypic evolution in dogs, if any, are unclear.

By using a variety of laboratory and analytical methods we have produced the first chromosomal-length assembly for the endangered Hawaiian monk seal. This study highlights how the improvements in sequencing technology have allowed even small groups to assemble high-quality genomes. This reference should be useful for continued studies of heterogeneity within the species and the consequences of inbreeding for future management. As is seen by the Vertebrate Genome Project, the DNAzoo, the Earth Biogenome Project and others the era of comparative genomics has arrived and the breadth of the data being produced is giving us important new perspectives on genome architecture and function.

## Acknowledgements

We thank Drs. Charles Littnan and Michelle Barbieri for obtaining samples, Dr. Susan Brown for performing the optical genome mapping for the first optical genome map, Dr. Melissa Olson for lymphocyte separation and whole blood cryopreservation, Laura Kasch and Jill Barton for preparing agarose blocks for first optical genome mapping experiment and DNA isolation from blood. Michael Campbell at 10X Genomics for assistance with Loupe viewer.

Hi-C scaffolding was performed by the DNA Zoo Consortium (www.dnazoo.org). DNA Zoo sequencing effort is supported by Illumina, Inc., IBM, and the Pawsey Supercomputing Center. E.L.A. was supported by the Welch Foundation (Q-1866), a McNair Medical Institute Scholar Award, an NIH Encyclopedia of DNA Elements Mapping Center Award (UM1HG009375), a US-Israel Binational Science Foundation Award (2019276), the Behavioral Plasticity Research Institute (NSF DBI-2021795), NSF Physics Frontiers Center Award (NSF PHY-2019745), and an NIH CEGS (RM1HG011016-01A1).

## Author Contributions

David Mohr: Sequencing, sequence assembly, methods development, manuscript text and figures

Justin Paschall: Illumina DRAGEN support for variants Ahmed Naguib: Bionano Genomics support

Andy Wing Chun Pang: Bionano Genomics support

Deanna Church: Genomics techniques advice and human reference

Stephen Gaughran: Population genetic analysis, WGS on additional animals, MSMC figure

Olga Dudchenko: Hi-C chromosome-length assembly and Juicebox Assembly Tools polishing

Erez Lieberman Aiden: DNA Zoo assembly effort supervision Alan Scott: Conceived project, manuscript text, figures

## Supplementary materials

**Supp. Table 1.** Methods used in this study and the order in which they were employed.

| Method | Platform | Input | Data type | Analysis tools |
| --- | --- | --- | --- | --- |
| DISCOVAR | Illumina HS2500 | 500ng sheared DNA sheared to ~400 bp | 2X250 PE reads | DISCOVAR *denovo* v52488 |
| 10X Genomics Chromium | Illumina HS2500 | 1.2ng HMW DNA (>40 kbp) | 2X150 PE reads | Supernova v1.1 And V2.+ |
| Optical genome mapping | Bionano Irys BspQ I | ~10<sup>7</sup> cells<br>~160ng HMW DNA | >150 kbp molecules | BNG Assembler v5122 |
| Oxford Nanopore | R9.1 Flow cell | 1-2 ug HMW DNA<br>~9X read depth | N50 ~32 kbp | LINKS, Pilon |
| Hi-C, Arima | Illumina NS6500 | ~1 M reads | 2X150 PE reads | 3d-dna, Juicebox tools |
| Optical genome map 2 | Saphyr DLE-1 | ~10<sup>7</sup> cells<br>~160ng HMW DNA | >150 kbp molecules | BNG Access |
| Synteny analysis |  |  | NCBI, DNazoo assemblies | Minimap2, D-GENIES |
| Manula editing | Galaxy tools |  |  |  |

**Supp. Table 2.** Assembly sources for species comparisons.

| Common name | Species | Assembly | Source |
| --- | --- | --- | --- |
| Gray Seal | *Halichoerus grypus* | Halichoerus_grypus_HiC | DNAZoo |
| Harbor Seal | *Phoca vitulina* | GSC_HSeal_1.0_HiC | DNAZoo |
| Bearded Seal | *Erignathus barbatus* | Erignathus_barbatus_HiC | DNAZoo |
| N. Elephant Seal | *Mirounga angustirostris* | Mirounga_angustirostris_HiC | DNAZoo |
| **Hawaiian Monk Seal** | ***Neomonachus schauinslandi*** | **GCF_002201575.2_ASM220157v2** | **NCBI** |
| Antarctic Fur Seal | *Arctocephalus gazella* | GCA_900642305.1_arcGaz3 | NCBI |
| Guadalupe Fur Seal | *Arctocephalus townsendi* |  |  |
| California Sea Lion | *Zalophus californianus* | GCA_900631625.1_zalCal2.2 | NCBI |
| Walrus | *Odobenus rosmarus divergens* | Oros_1.0_HiC | DNAZoo |
| Domestic cat | *Felis catus* | GCF_000181335.3_Felis_catus_9.0 | NCBI |
| Domestic dog | *Canis familiaris* | GCF_000002285.3_CanFam3.1 | NCBI |

**Supp. Table 3.** Comparison of NCBI metrics for ASM220157v1 ASM220157v2.

| NCBI Global statistic | V1 | V2 |
| --- | --- | --- |
| Total ungapped length | 2,347,217,317 | 2,376,537,768 |
| NCBI annotations |  |  |
| Genes and Pseudogenes | 24,730 | 27,736 |
| Protein coding genes | 18,856 | 19,838 |
| Exons | 204,392 | 212,563 |
| Non-coding | 1,173 | 3,122 |
| Pseudogenes | 4,701 | 4,734 |
| mRNAs | 28,368 | 34,516 |
| tRNAs | 446 | 531 |
| lncRNA | 824 | 780 |

**Supp. Fig. 1.**
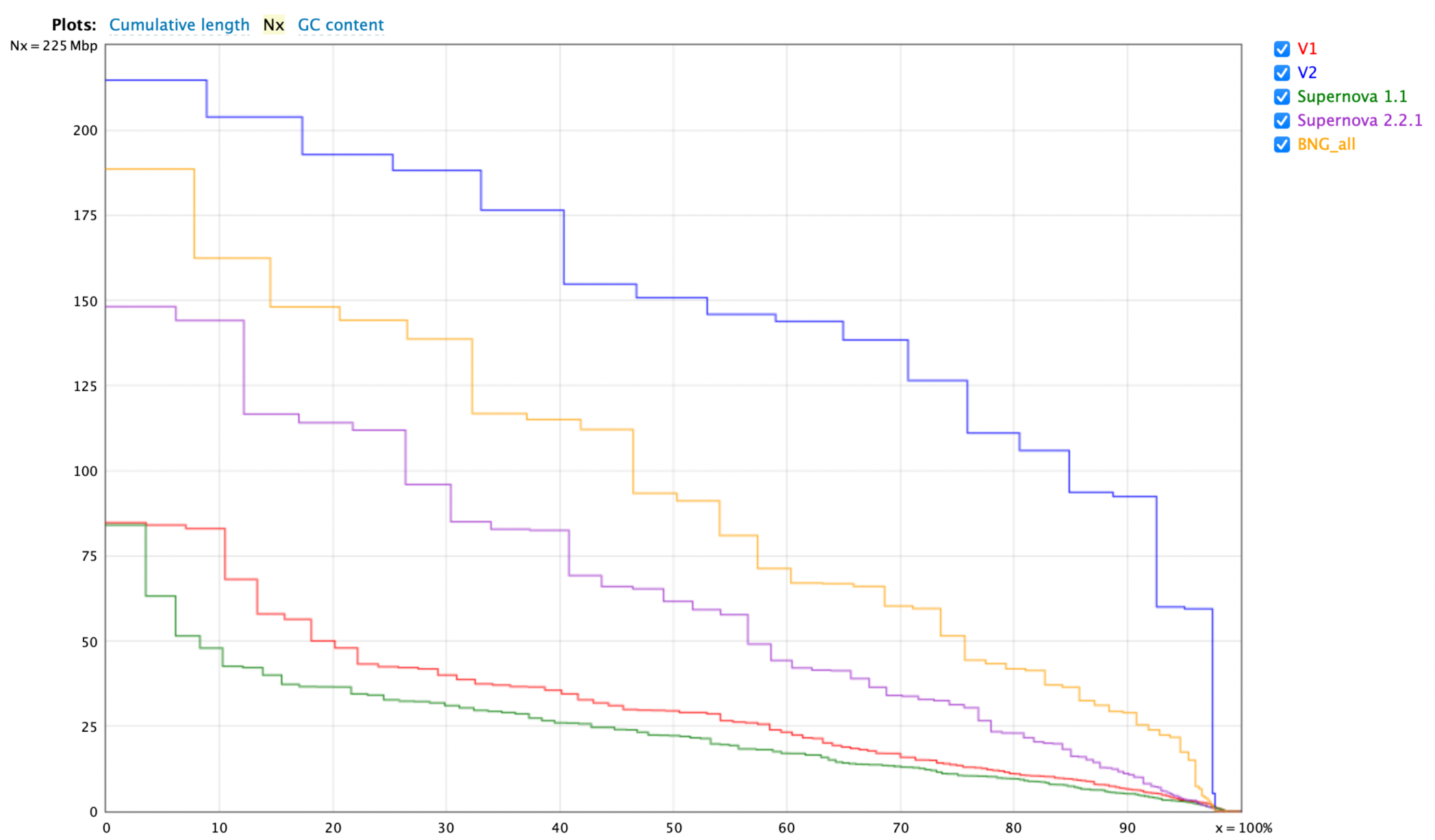
Improved assembly contiguity with additional methods. Linked reads with Supernova v1.1, N50=22.23 Mb; ASM220157v1 assembly, N50=29.52 Mb; Supernova 2.2.1 on original linked reads, N50=61.65 Mb; All methods with BNG maps, N50=93.39 Mb; ASM220157v2 assembly, 150.81 Mb.

**Supp. Fig. 2.**
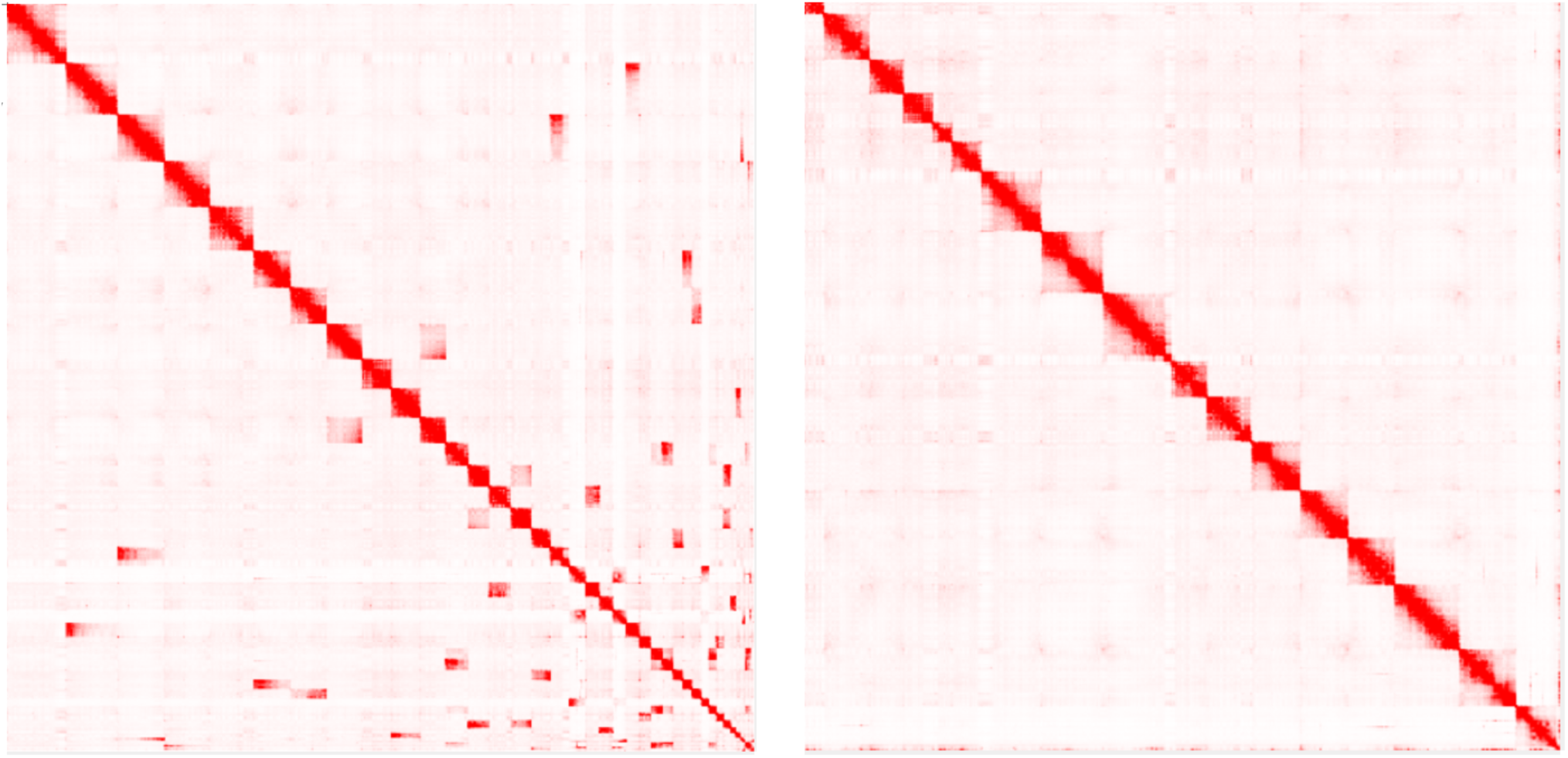
Hi-C assembly before and after manual editing with Juicer. Original plots and data are available at DNAzoo.org.

**Supp. Fig. 3.**
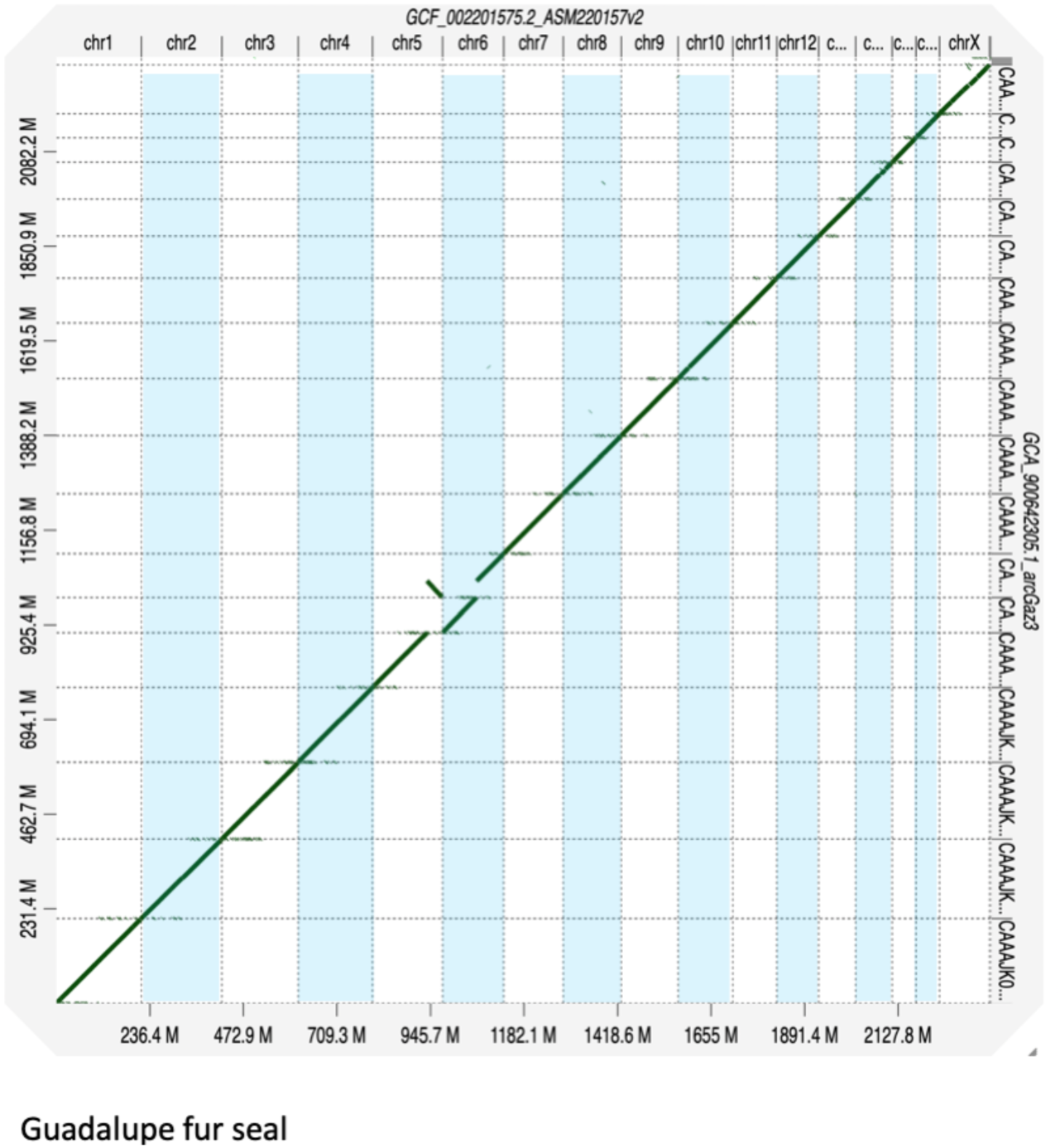
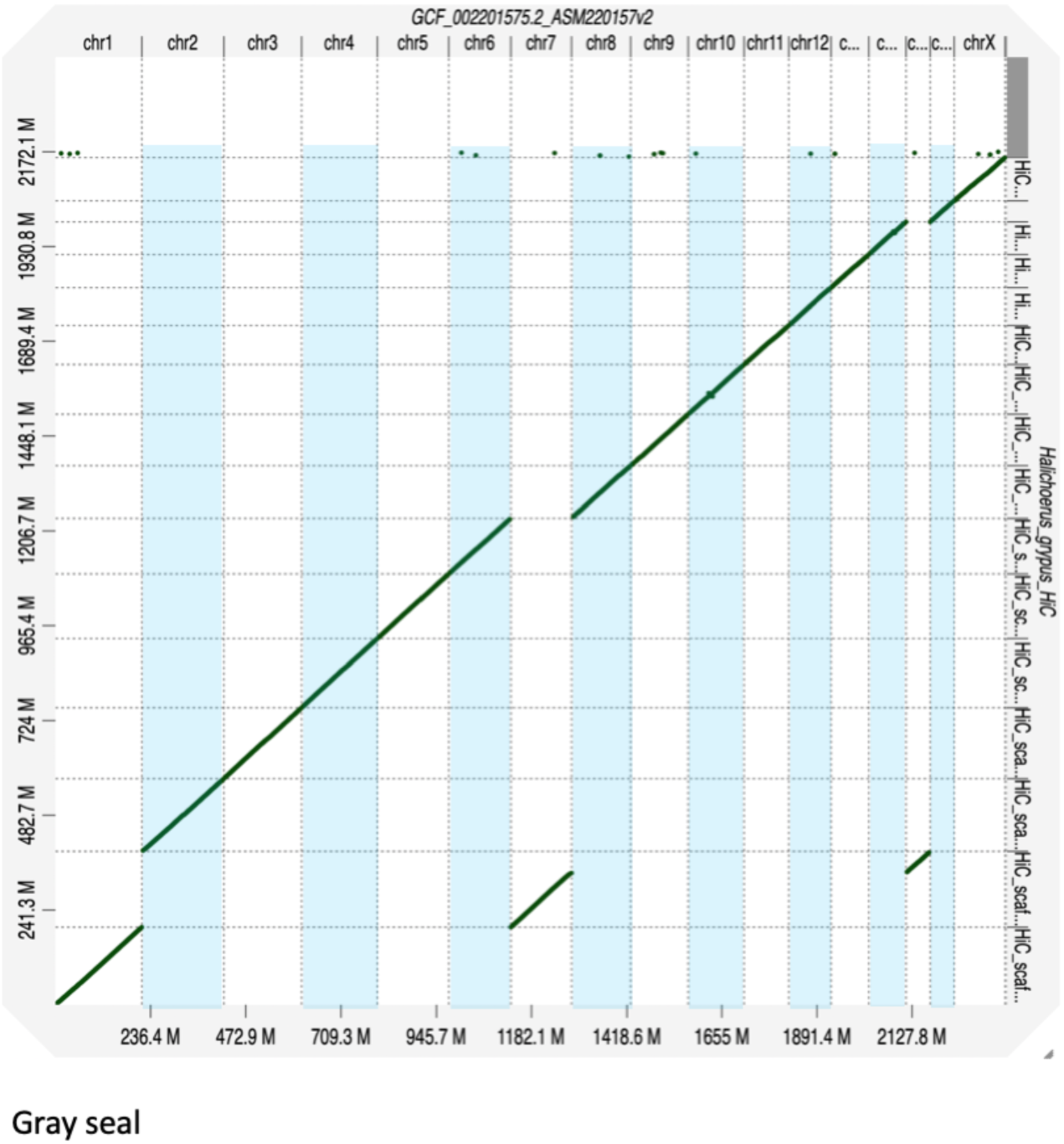
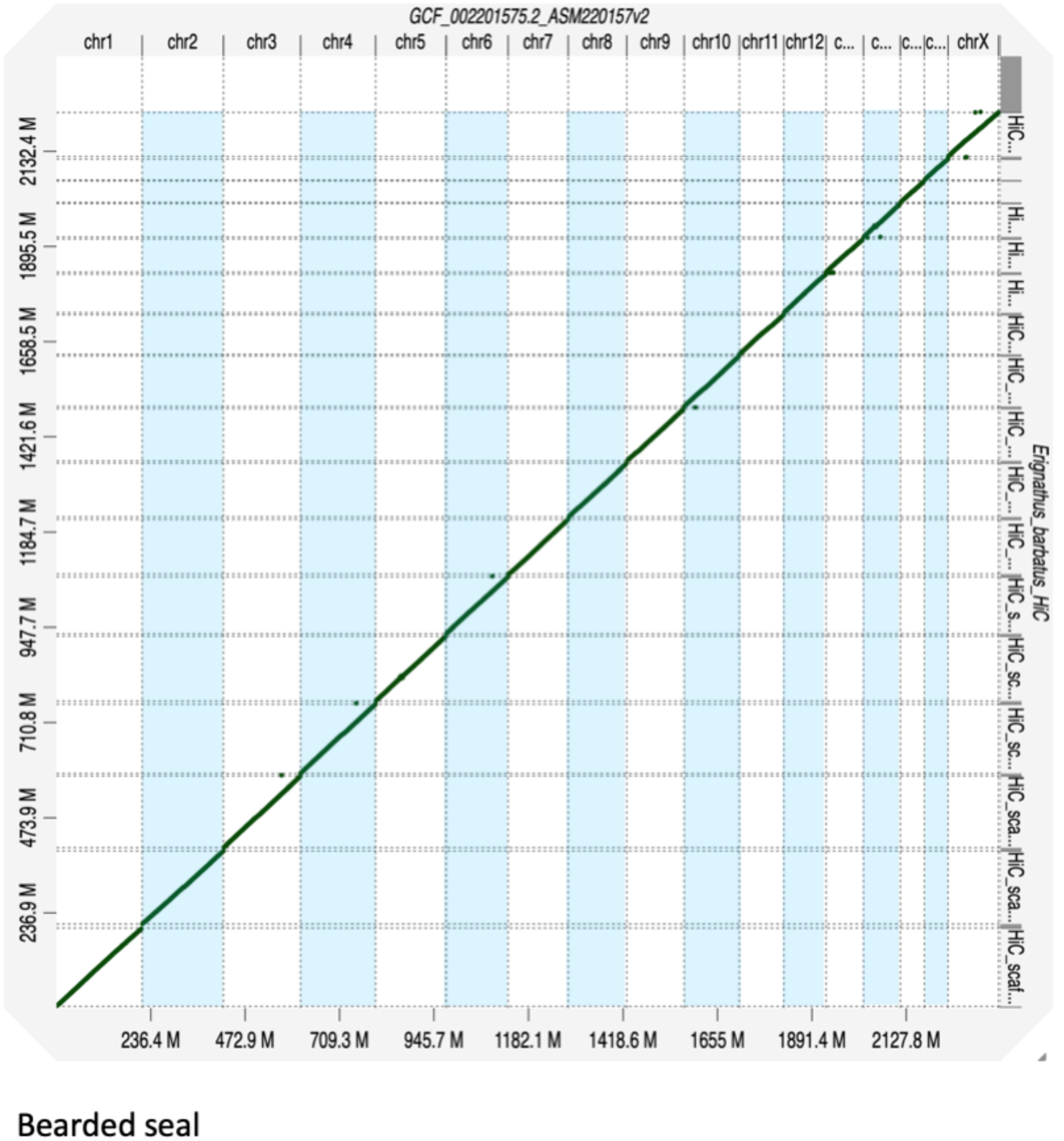
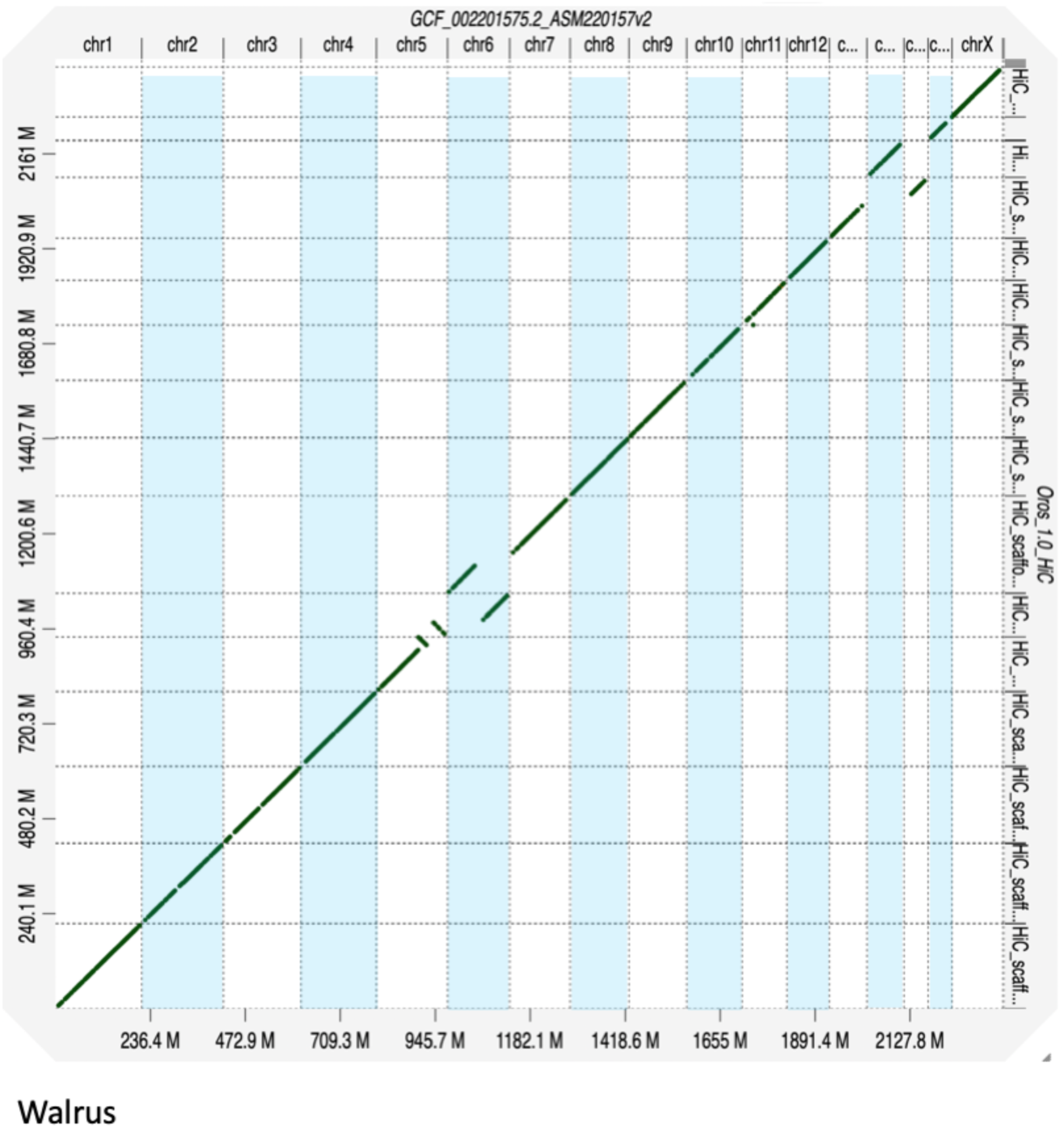
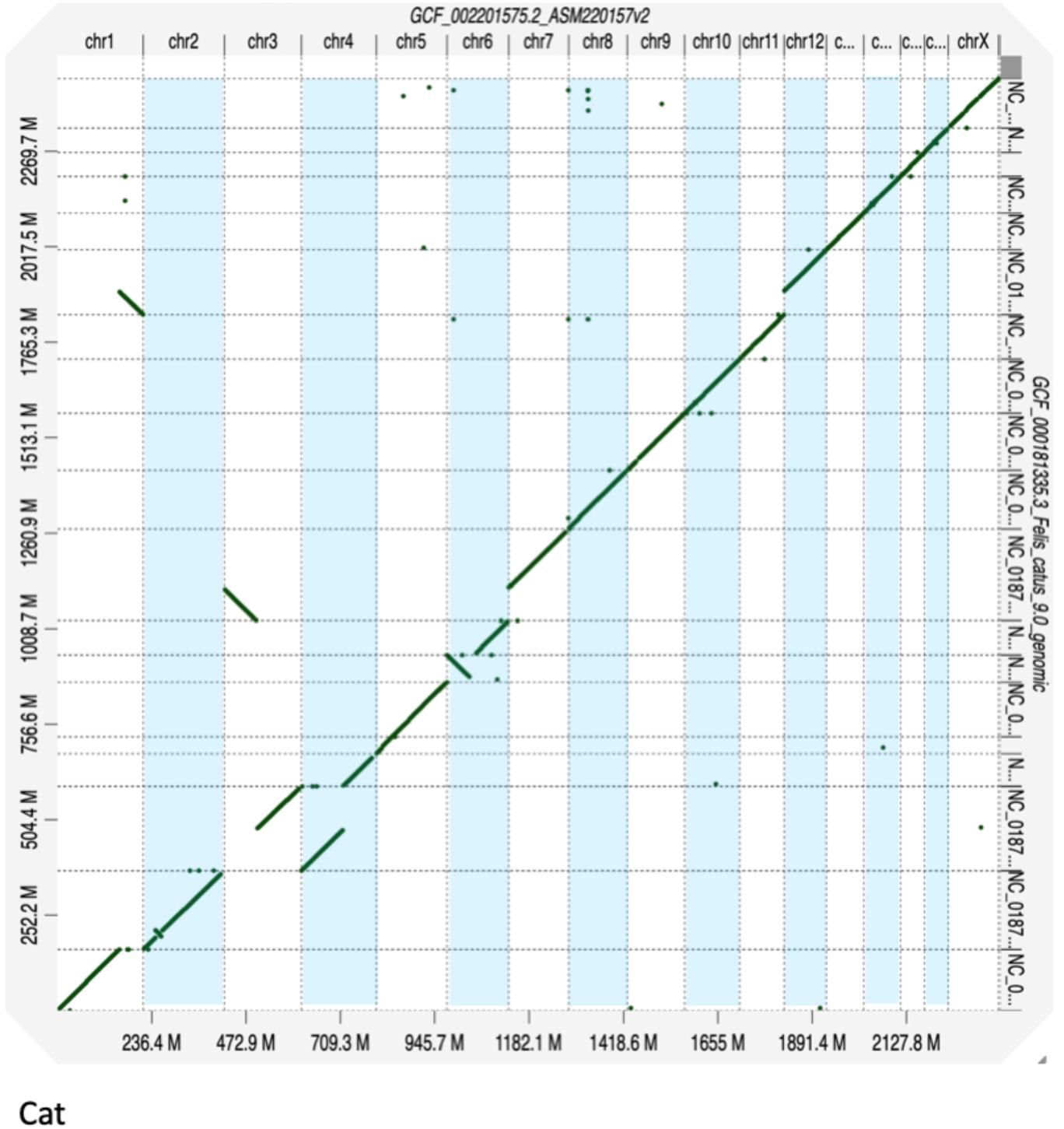
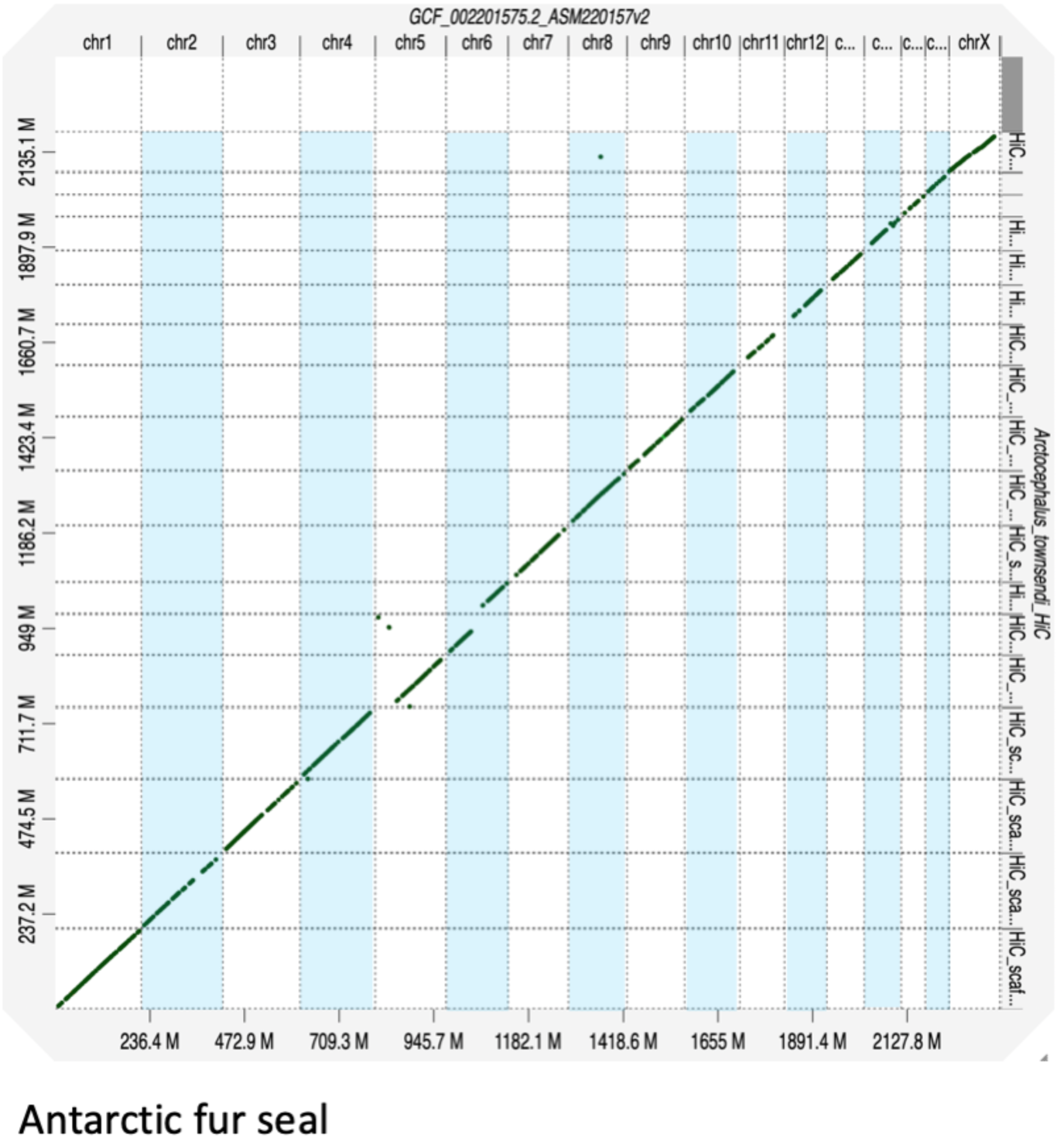
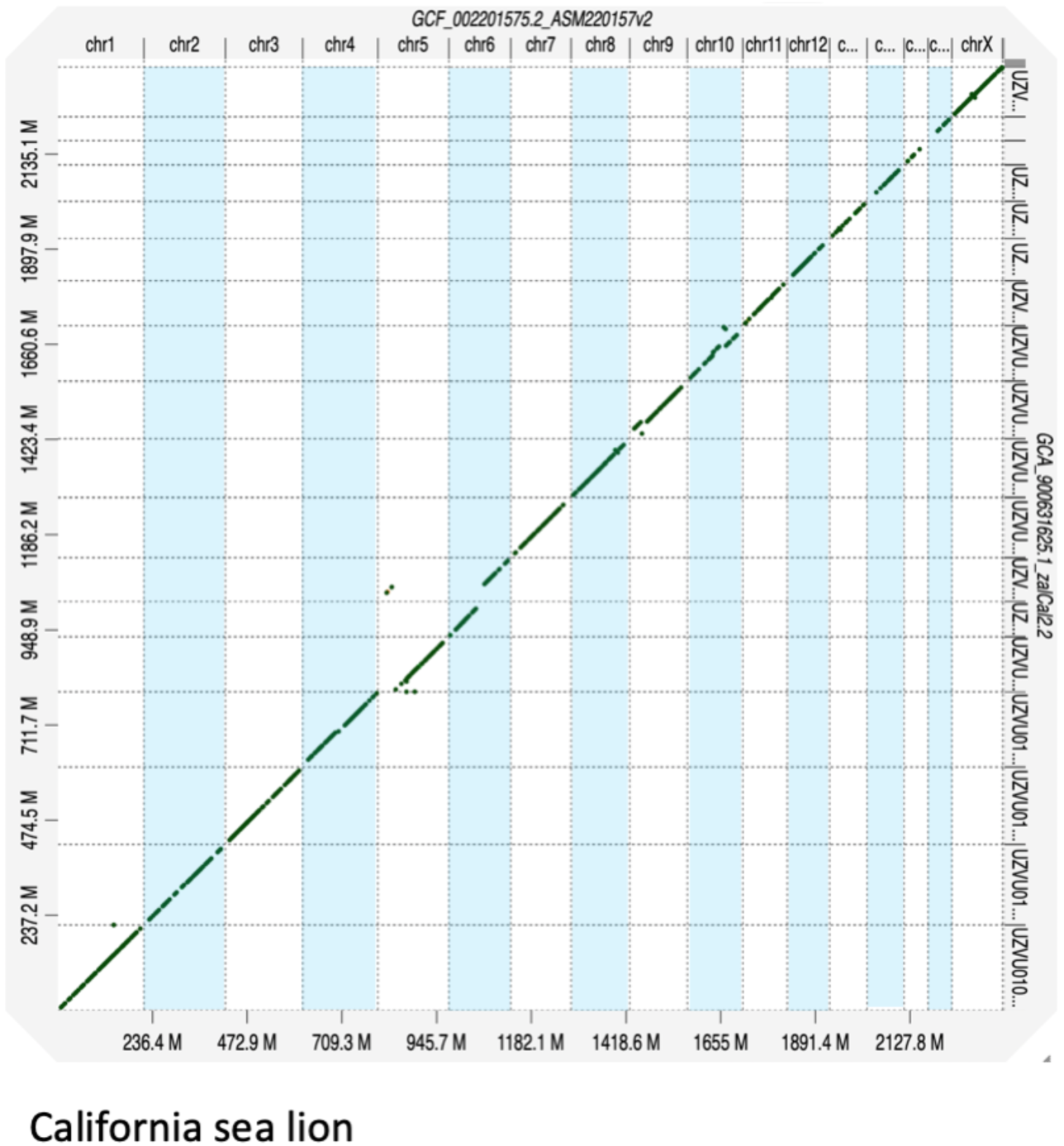
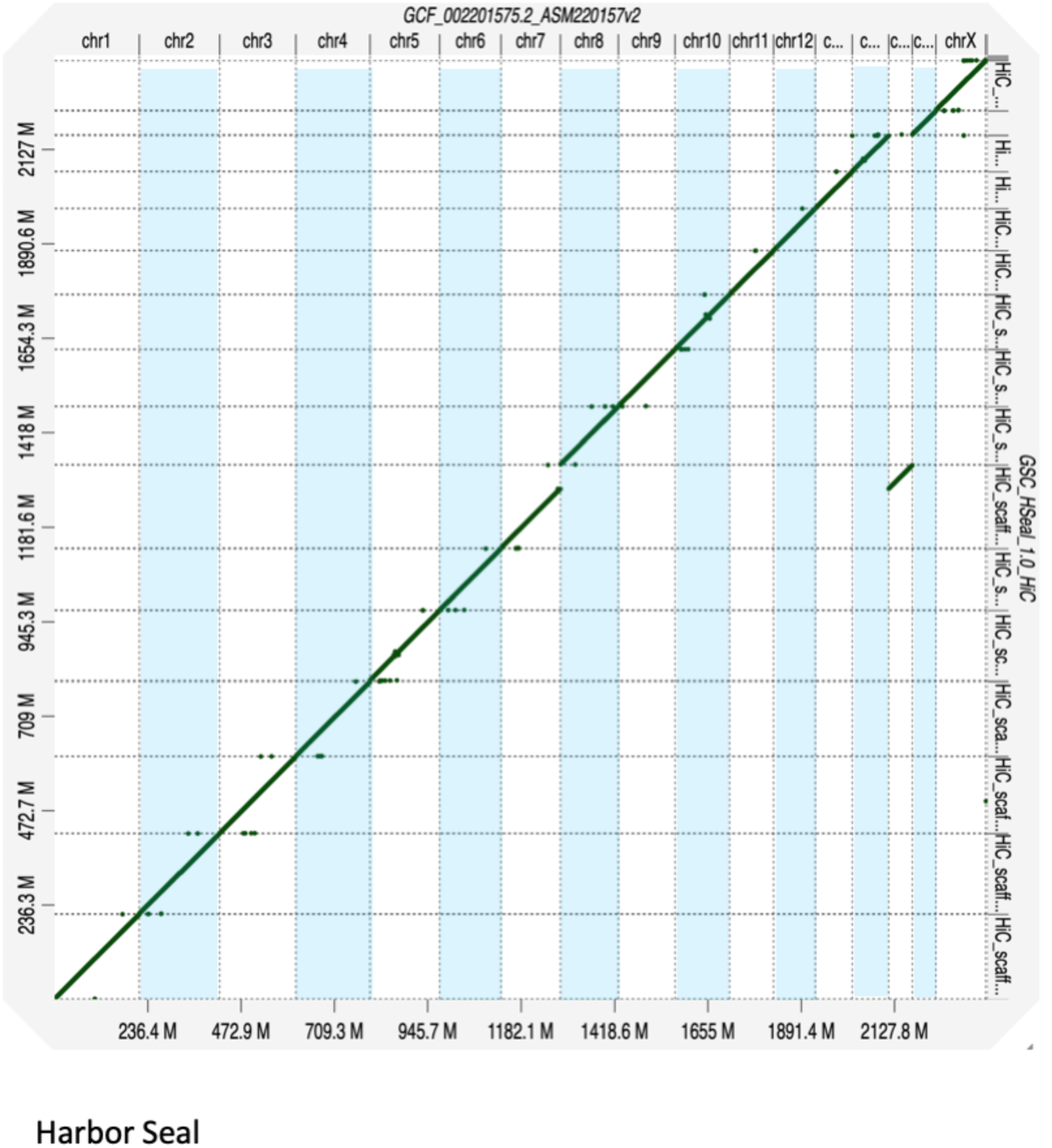
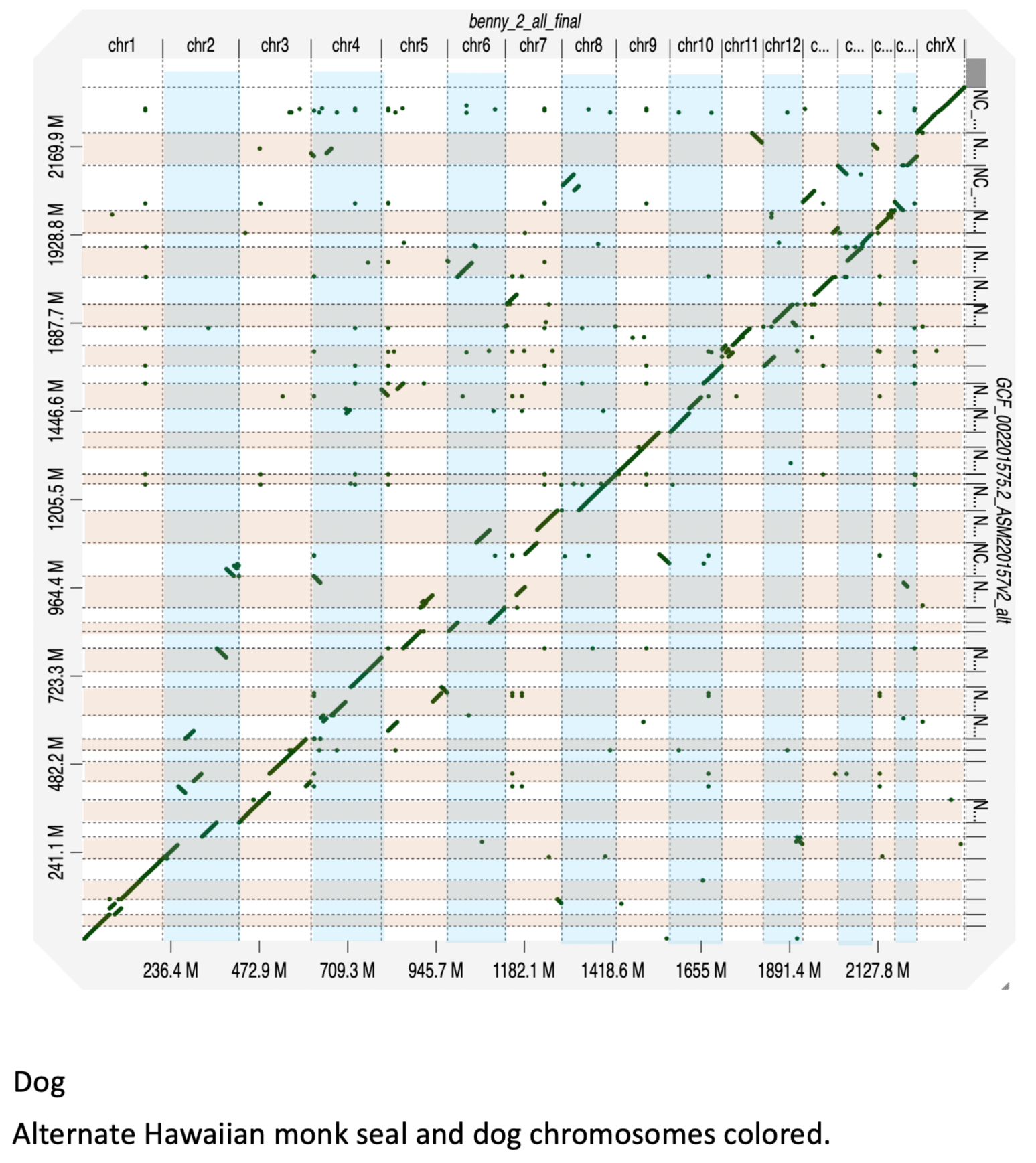
Enlarged DG plots of other species relative to HMS ASM220157v2. Selected chromosomes from other species were reverse-complemented before plotting. [[DRAFT FIG.]]

**Supp. Fig. 4.**
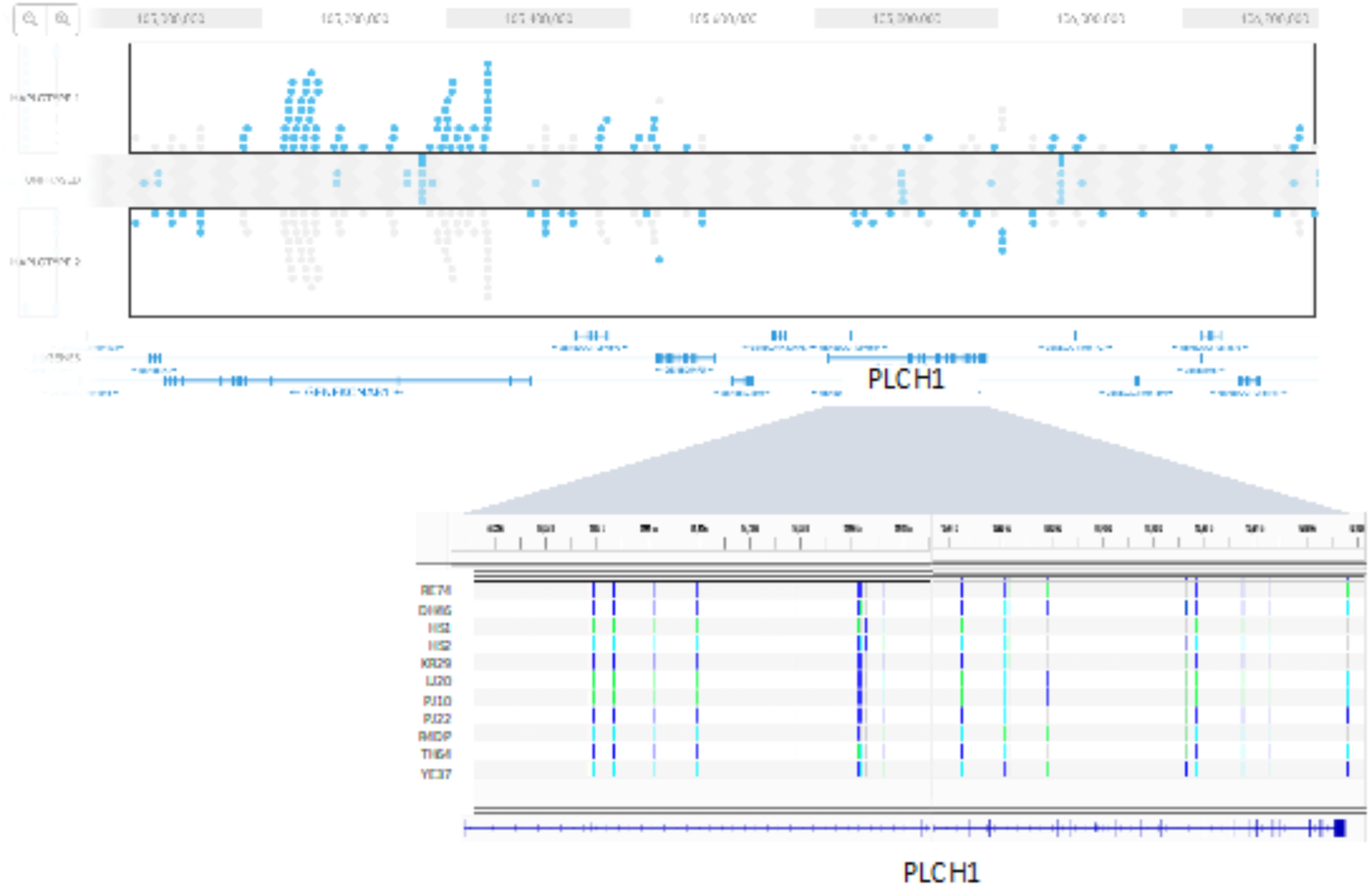
A representative 1.28Mb phase block in RE74 including KCNAB1 and PLCH1 shown in the Loupe viewer. Alignment of variants in RE74 and 10 other seals within PLCH1. Solid dark blue bars are heterozygous differences from the RE74 reference, light blue are homozygous differences from reference and green bars are homozygous reference.

## Footnotes

1 M. schauinslandi was moved to the genus Neomonachus in 2014 (Scheel et al, 2014)

